# The role of N-glycans and their processing in ER-to-lysosome-associated degradation of disease-causing mutant Neuroserpin

**DOI:** 10.64898/2026.04.02.716018

**Authors:** Carolin Hoefner, Ilaria Fregno, Maurizio Molinari

## Abstract

Most proteins synthesized in the endoplasmic reticulum (ER) are covalently modified upon addition of pre-assembled oligosaccharides to side chains of asparagine (N) residues. Processing of N-linked oligosaccharides by ER-resident glucosidases, mannosidases and glucosyltransferases determines the fate of the associated polypeptides. Terminally glucose residues are removed from N-glycans to hamper engagement of ER-resident glucose-binding chaperones and promote secretion of native polypeptides. Mannose residues are removed to target terminally misfolded proteins for dislocation across the ER membrane and clearance by the cytoplasmic ubiquitin proteasome system (ER-associated degradation, ERAD). Recent evidence highlights the role of persistent N-glycan glucosylation as a signal that promotes segregation of misfolded proteins in ER subdomains that are eventually delivered to endolysosomal compartments for ER-to-Lysosome-Associated Degradation (ERLAD). Here we show that the polymerization-prone Portland variant of Neuroserpin (NS_PL) associated with familial encephalopathy with NS inclusion bodies (FENIB) is a client of the ERLAD machinery. Its lysosomal clearance relies on the LC3-dependent delivery branch of ERLAD involving the lectin chaperone Calnexin (CNX), the ERphagy receptor FAM134B and the SNARE protein Syntaxin17 (STX17), which is engaged upon persistent glucosylation of the NS_PL oligosaccharide linked at the asparagine residue at position 321.

## Introduc1on

The ER is the main biosynthetic organelle of nucleated cells, producing lipids, oligosaccharides, and a large fraction of the cellular proteome [1]. Most proteins synthesized in the ER are modified by N-linked glycosylation. N-glycosylation is a critical post-translational modification in eukaryotes and archaea, where a pre-assembled sugar chain (glycan) is attached to the side chain of an asparagine residue in an asparagine-any amino acid-serine/threonine (Asn-Xxx-Ser/Thr) consensus sequence. N-glycosylation is operated co- or post-translationally by the oligosaccharyltransferase (OST) complex associated with the protein translocation machinery at the ER membrane [2]. N-glycans are composed of 3 glucose, 9 mannose and 3 N-acetylglucosamine residues. Their sequential processing by ER-resident glucosidases, mannosidases and glucosyltransferases determines the fate of the associated polypeptide, which is either retained in the ER to complete the folding program, secreted upon achievement of the native structure, or eventually degraded if folding fails [3]. Despite a folding program assisted by high concentrations of molecular chaperones and folding enzymes, up to one third of newly synthesized proteins fail to reach the native state [4]. To maintain a healthy proteome and ensure cellular homeostasis, terminally misfolded proteins are efficiently eliminated either by ER-associated degradation (ERAD) or by ER-to-Lysosome-Associated Degradation (ERLAD) [5–8]. Engagement of the ERAD machinery relies on slow removal of terminal mannose residues from N-linked glycans of misfolded polypeptides that facilitates retro-translocation across the ER membrane, polyubiquitylation, and subsequent proteasomal degradation [9–11]. Misfolded proteins that fail to enter the ERAD pathways, enter instead ERLAD programs. These are initiated by glucose processing of misfolded proteins’ N-linked glycans that prolongs the association with ER-resident lectin chaperones, followed by segregation into specialized ER subdomains for lysosomal removal [12,13]. The delivery of misfolded polypeptides from the ER to the degradative compartments via ERLAD proceeds through three distinct routes: (i) macro-ERphagy, in which the ER subdomains containing misfolded polypeptides are engulfed by autophagosomes that eventually fuse with endolysosomes to generate the degradative autolysosomes [14]; (ii) micro-ERphagy, involving direct capture of ER subdomains by endolysosomes [15,16]; (iii) LC3-mediated vesicular transport, where ER-derived vesicles containing the misfolded proteins to be removed from cells fuse directly with lysosomes [5].

Neuroserpin (NS) is a neuronal serine protease inhibitor that regulates tissue plasminogen activator activity and supports synaptic function [17,18]. This soluble glycoprotein is synthesized in the ER of neurons. After proper folding, NS is secreted from the ER and is released in the extracellular space through the Golgi apparatus to axons and synapses [19]. Mutations in the SERPINI1 gene may introduce amino acid mutations in the NS polypeptide that destabilize the serpin fold, may promote aggregates formation and, if aggregates’ clearance is inefficient, may result in intraluminal retention of the mutant polypeptide in soluble or insoluble form. This is linked to a hereditary neurodegenerative disease called familial encephalopathy with NS inclusion bodies (FENIB) [19,20]. Here, we investigate the intracellular fate of the Portland mutation of NS (NS_PL). NS_PL is characterized by the replacement of the serine (S) residue at position 52 in the polypeptide sequence with an arginine (R) (S52R, pink in **Fig. 1A**). NS_PL is one of the aggregation-prone NS variants associated with FENIB [17,20–22]. Previous studies showed that clearance from the ER of this folding-defective polypeptide relies on the intervention of both the ubiquitin-proteasome system and autophagy. The autophagic clearance of NS_PL was pharmacologically inhibited in cells exposed to Bafilomycin A1 (BafA1) to inactivate lysosomal hydrolases and by genetic inactivation of autophagy upon deletion of ATG5 [25]. We first confirmed these data to then establish in molecular detail the lysosomal clearance pathway that ensure the removal of NS_PL aggregates from the ER lumen. Our study shows that terminally misfolded NS_PL is an ERLAD client and reveals that lysosomal delivery of this polypeptide requires glucose processing of the N-glycan at position 321 and involves the lectin chaperone CNX, the ERphagy receptor FAM134B, the SNARE protein STX17 and is dependent on LC3 lipidation.

**Fig. 1.**
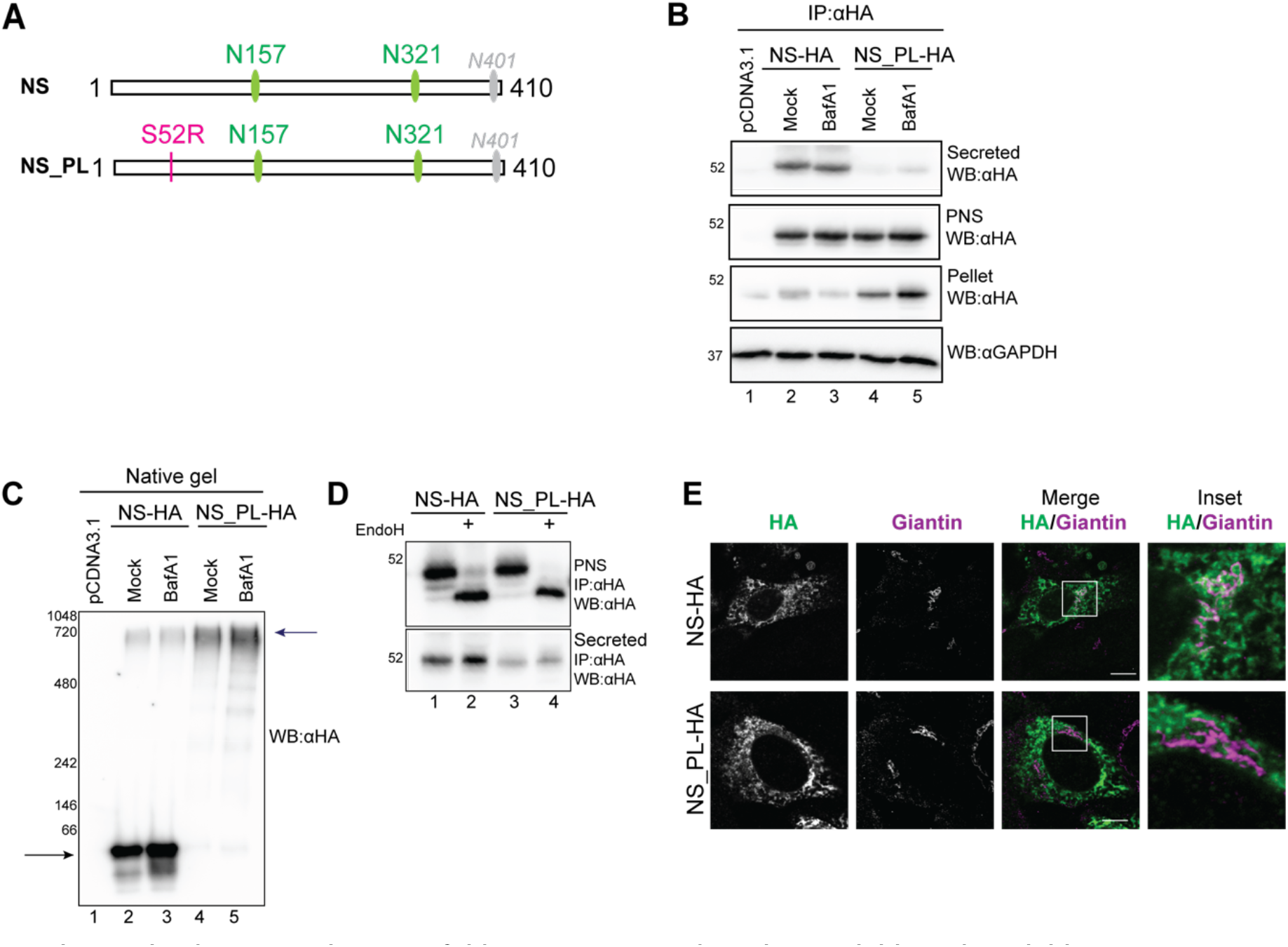
The Portland mutation hampers folding, secretion and results in soluble and insoluble aggregates. **A)** The three potential N-glycosylation sites of NS and NS_PL are shown in the schematics. Numbers refer to the position of the asparagine residue (N)-acceptor of the oligosaccharide. The N401 site is not glycosylated. The S52R mutation of NS_PL is shown. **B)** HEK293 cells were transiently transfected with NS-HA or NS_PL-HA and mock-treated or exposed to 100nM BafA1. NS-HA and NS_PL-HA were immunoisolated with anti-HA antibodies from distinct cellular fractions (secreted, detergent-soluble, and detergent-insoluble). Proteins were separated in SDS-PAGE and revealed by immunoblotting. **C)** Same as (B) where proteins in the detergent-soluble intracellular fraction were separated in native gels. Monomers (lanes 2, 3) and polymers (lanes 4, 5) are shown with arrows. **D)** EndoH sensitivity assay of the glycoproteins immunoisolated from the cell media or the cytoplasmic fraction. **E)** CLSM analyses in WT MEF. NS-HA and NS_PL-HA (green) with the Golgi marker Giantin (magenta). Scale bars 10 µm.

## Results

### The Portland mutation enhances NS misfolding and deposition in insoluble aggregates

Firstly, we monitored the intracellular fate of HA-tagged versions of NS or NS_PL transiently expressed in Human Embryonic Kidney HEK293 cells. Twelve hours after transfection, the medium was replaced with treatment-containing medium, and the supernatant was collected 6 h later for analysis, whereas the cells were detergent-solubilized as described in the Methods section. The presence of NS and NS_PL was monitored by immunoblotting in the cell culture media (**Fig. 1B**, upper panel, secreted), in the cell lysate (**Fig. 1B**, second panel, post nuclear supernatant (PNS)) and in the detergent-insoluble fraction (**Fig. 1B**, third panel, pellet). The mutation in NS_PL substantially inhibited the secretion of the ectopically expressed polypeptide (**Fig. 1B**, upper panel, lane 4) compared to the folding competent NS (**Fig. 1B**, upper panel, lane 2). In the detergent-soluble intracellular fraction, we detected comparable abundance for both variants (**Fig. 1B**, second panel, lanes 2 *vs*. 4). However, analysis of these fractions in native gels revealed that the NS was present in the detergent soluble fraction as a monomer (arrows, **Fig. 1C**, lane 2), whereas the NS_PL was in high molecular weight complexes, as expected from its propensity to undergo polymerization [23] (arrows, **Fig. 1C**, lane 4). Also, and in sharp contrast to NS (**Fig. 1B**, third panel, lanes 2-3), NS_PL showed a marked accumulation in the detergent-insoluble fraction (**Fig. 1B**, third panel, lane 4).

To confirm that the NS and NS_PL harvested from the cell culture media was actively secreted from cells (and not released from dead cells), we performed an endoglycanase H (EndoH) assay. EndoH cleaves N-glycans of ER-localized glycoproteins but fails to cleave the glycans of polypeptides actively secreted from cells via the Golgi complex. These protein-bound oligosaccharides are in fact processed by Golgi enzymes to become EndoH-resistant [24]. When subjected to the EndoH assay, both the NS and NS_PL harvested from the cell culture media did not change their electrophoretic mobility. Thus, their N-glycans are EndoH-resistant testifying the secretion of the glycoproteins via the Golgi complex (**Fig. 1D**, upper panel, lanes 1 *vs*. 2 for NS and lanes 3 *vs*. 4 for NS_PL). In contrast, the polypeptides immune-isolated from the cells’ lysate were EndoH-sensitive and showed enhanced electrophoretic mobility upon EndoH treatment (**Fig. 1D**, second panel, lanes 1 *vs*. 2 and 3 *vs*. 4). Notably, a small fraction of intracellular NS was EndoH-resistant (**Fig. 1D**, second panel, lanes 1 *vs*. 2), consistent with the transit of this polypeptide in the Golgi Complex during secretion as testified by its co-localization with the Golgi marker Giantin in confocal laser scanning microscopy (CLSM, **Fig. 1E**, upper panel). Confirming previous data showing that autophagic pathways are involved in the luminal clearance of NS_PL aggregates [25], both the NS_PL in soluble (**Fig. 1C**, lanes 4 *vs*. 5) and in insoluble aggregates (**Fig. 1B**, third panel, lanes 4 *vs*. 5) accumulated upon inhibition of lysosomal activity in cells exposed to BafA1.

### NS_PL is delivered to LAMP1-positive endolysosomes for clearance

The propensity of misfolded NS_PL to form high order aggregates (**Fig. 1C**, lanes 4-5) and the accumulation of the protein in the insoluble fraction upon inactivation of the lysosomal activity in cells exposed to BafA1 (**Fig. 1B**, third panel, lanes 4-5) led us to check whether NS_PL is delivered within LAMP1-positive degradative endolysosomes for clearance. To this end, the subcellular distribution of NS_PL was examined in wild type mouse embryonic fibroblasts (MEF) in the absence (**Fig. 2A**) or in the presence (**Fig. 2B**) of BafA1, which inhibits lysosomal hydrolases thus resulting in accumulation of lysosomal clients in the lumen of LAMP1-positive degradative compartments [14,26]. In cells with functional endolysosomes, NS_PL was not found within the LAMP1-positive compartments (**Fig. 2A**). The substantial accumulation of NS_PL within the LAMP1-positive endolysosomes upon inactivation of their degradative activity in cells exposed to BafA1 (**Fig. 2B**, quantification in **Fig. 2H**) confirmed that NS_PL is transported from the ER to endolysosomes for clearance and identifies NS_PL as a possible client of the ERLAD machinery.

**Fig. 2.**
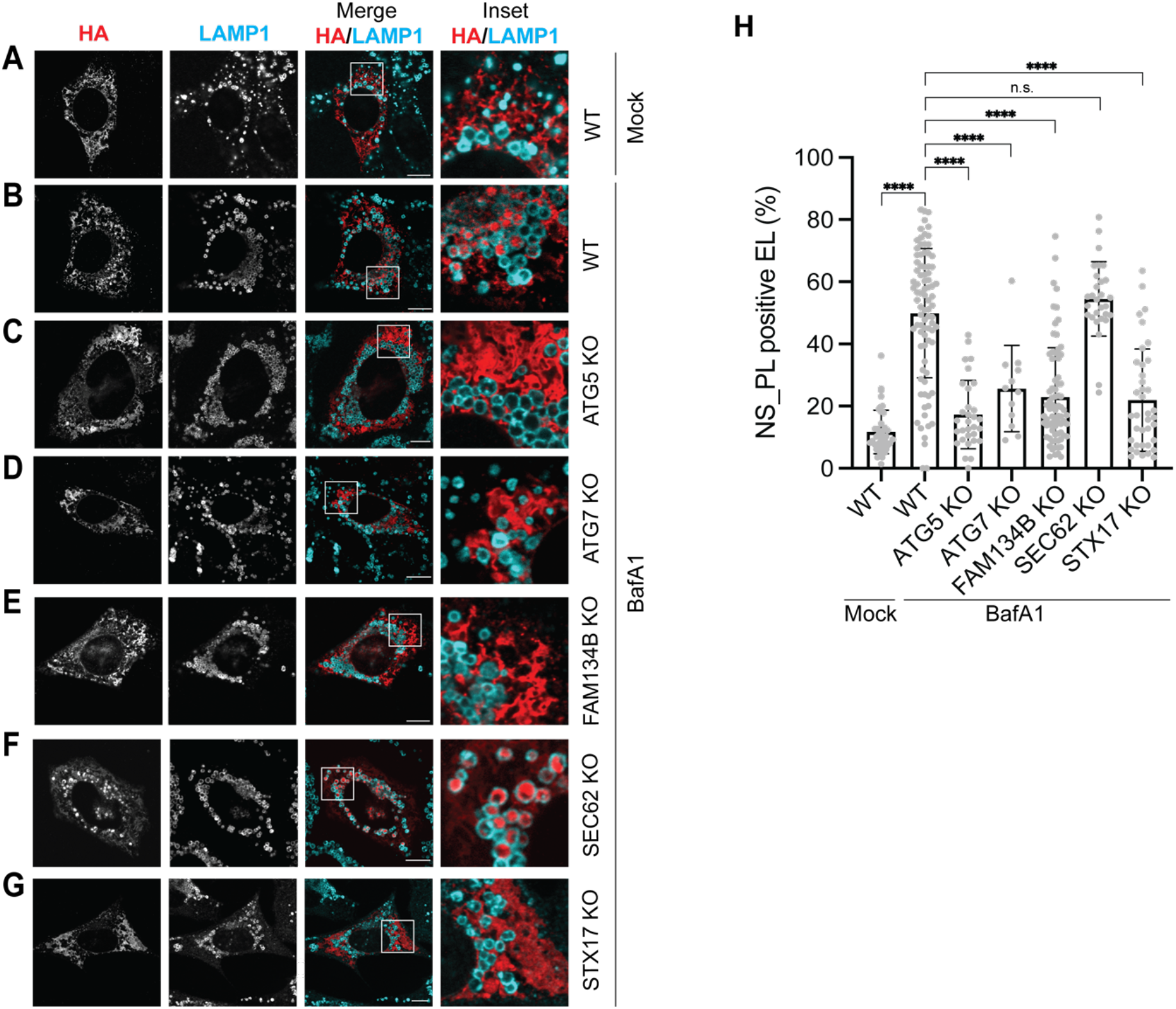
Monitoring the delivery of NS_PL to LAMP1-positive endolysosomes for degradation. **A)** CLSM analyses in WT MEF. NS_PL-HA (red) in LAMP1-positive endolysosomes (cyan). Scale bars 10 µm. **B)** Same as (A) in MEF treated with BafA1. **C)** Same as (B) in ATG5 KO MEF. **D)** Same as (B) in ATG7 KO MEF. **E)** Same as (B) in FAM134B KO MEF. **F)** Same as (B) in SEC62 KO MEF. **G)** Same as (B) in STX17 KO MEF. **H)** LysoQuant [28] quantification of HA-positive endolysosomes of A-G (N=3 for C, D, F, G; N=5 for A, B, E). Mean, n = 41, 85, 31, 12, 74, 29, 38 cells. Welch’s T test of One-way ANOVA, ns P > 0.05, ****P < 0.0001.

### NS_PL is an ERLAD client

Next, we monitored by CLSM the delivery of NS_PL in the LAMP1-positive degradative compartment in a series of cell lines characterized by defective LC3 lipidation and dysfunctional autophagy (ATG5- and ATG7-KO), lacking the ERphagy receptors FAM134B or SEC62, or lacking the SNARE protein STX17, which controls membrane:membrane fusion events between the ER and degradative endolysosomes (**Fig. 2**) [12,13,27].

LC3 lipidation is a defining step in selective autophagy pathways as it attaches LC3 to membranes and thereby allows autophagy receptors to link cargos to the forming autophagosome or to degradative endolysosomes [14,29]. To assess whether lysosomal delivery of NS_PL depends on the LC3 lipidation machinery, the folding-defective polypeptide was transiently expressed in MEF lacking ATG5 (**Fig. 2C**) [30] or ATG7 (**Fig. 2D**) [31], two essential components required for LC3 conjugation. Loss of either ATG5 or ATG7 markedly reduced the delivery of NS_PL to endolysosomes (**Figs. 2C-2D**, respectively, **Fig. 2H**).

Next, to determine whether an ERphagy receptor mediates the delivery of NS_PL to endolysosomes for clearance, the folding-defective polypeptide was transiently expressed in MEF lacking the ERphagy receptors FAM134B (**Fig. 2E**, CRISPR cells are described in [12]), or SEC62 (**Fig. 2F**, CRISPR cells are described in [32]). We observed that deletion of FAM134B substantially inhibited endolysosomal delivery of NS_PL (**Figs. 2E, 2H**), whereas deletion of SEC62 had no consequence (**Figs. 2F, 2H**). Finally, the substantial decrease of delivery of NS_PL to endolysosomes in STX17 KO MEF (**Figs. 2G, 2H** CRISPR cells are described in [12]) confirmed that fusion events between the ER and the endolysosomal membrane are required to deliver the ERLAD client from the ER lumen to the endolysosomal lumen. Taken together, these data reveal that NS_PL is a canonical client of the ERLAD machinery [6].

### N-glycan processing channels NS_PL in the ERLAD machinery

The disease-causing NS_PL variant of NS is a di-glycosylated protein (the cryptic glycosylation site at position 401 is not modified in the NS and NS_PL protein, **Fig. 1A**) [22].

N-linked glycans are rapidly processed by the α_1,2_-glucosidase I (GI in **Fig. 3A**) that removes the outermost glucose 1, and by the α_1,2_-glucosidase II (GII) that removes glucose 2. These cleavages expose the innermost glucose 3 (circled in red in **Fig. 3A**) which engages the lectin-like chaperone CNX [2]. GII also removes the innermost glucose 3 to disengage CNX. If the folding program has been completed, the mature polypeptide is released in the secretory line to be secreted from cells. If the protein is still unfolded, as it is the case for mutant proteins like NS_PL, the UDP-glucose:glycoprotein glucosyltransferase (UGGT1 in **Fig. 3A**) reglucosylates the N-linked oligosaccharides to prolong CNX engagement [2]. We previously reported in the case of the Z-variant of α1 antitrypsin that the sustained interaction with CNX drives the segregation of terminally misfolded polypeptides into ER subdomains enriched in the ERphagy receptor FAM134B [12,13,33]. To assess whether the clearance of NS_PL from the ER follows this route, the lysosomal delivery of the folding-defective protein was assessed by CLSM in MEF mock treated or subjected to pharmacologic inactivation of glycoprotein association with CNX. This was achieved by incubating cells with Castanospermine (CST), a cell permeable inhibitor of ER-resident GI and GII [2,11,34]. As shown in **Fig. 2**, NS_PL accumulated within the LAMP1-positive compartments in MEF exposed to BafA1 (**Figs. 3B** *vs*. **3C, 3G**). Cells exposure to CST drastically reduced the lysosomal delivery of NS_PL (**Figs. 3D, 3G**). Likewise, genetic inactivation of the CNX chaperone system upon deletion of the glucosylation enzyme UGGT1 [35,36] or upon deletion of CNX [37–39] substantially inhibited lysosomal delivery of NS_PL (**Figs. 3E-3F, respectively and 3G**). Altogether these results show the importance of glucose processing and persistent lectin-engagement in FAM134B-driven NS_PL selection for ERLAD.

**Fig. 3.**
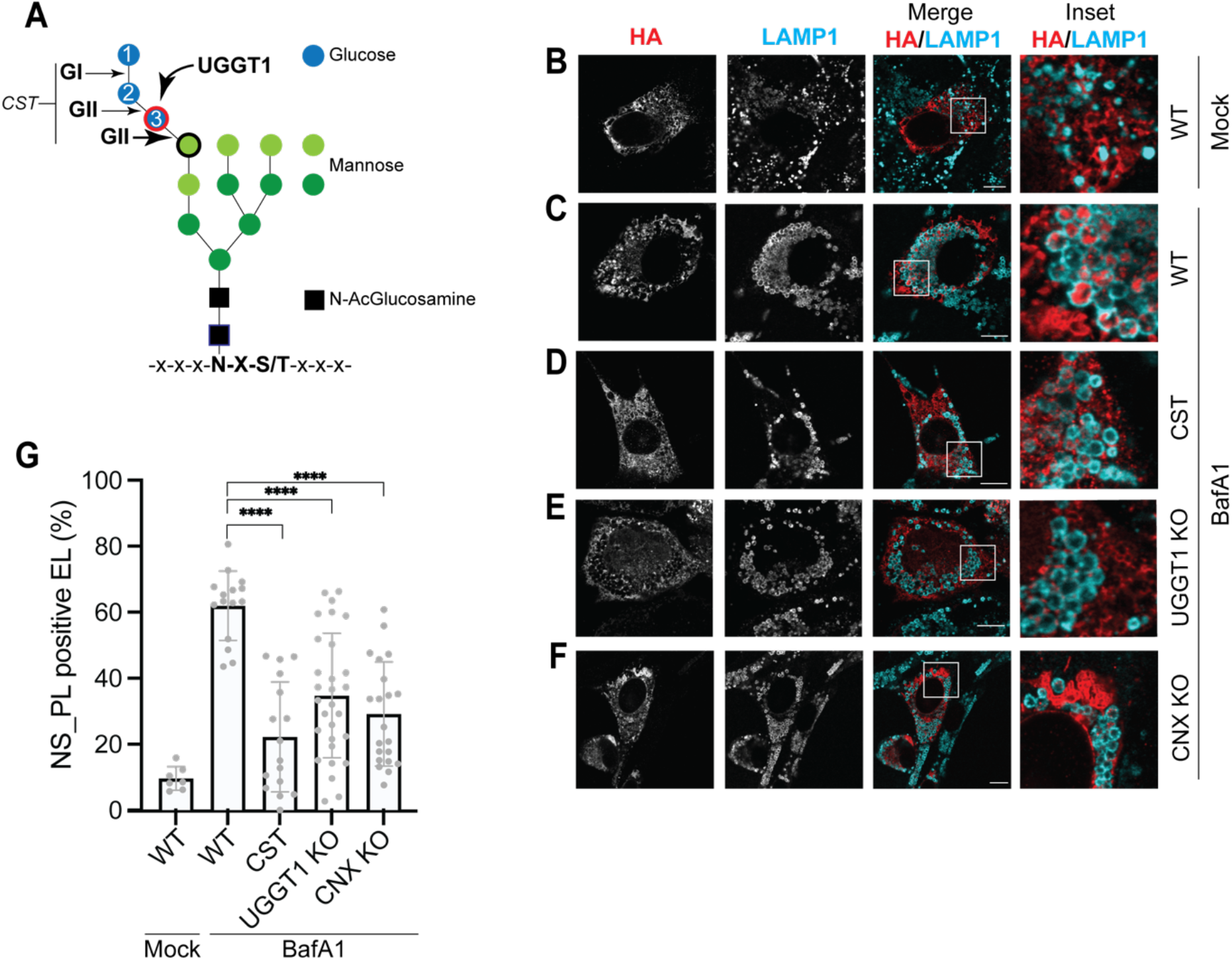
CNX and glucose processing enzymes regulate NS_PL delivery to endolysosomes. **A)** N-glycan. Glucoses, blue; mannoses, green; N-acetylglucosamines, black. GI removes glucose 1. GII removes glucose residues 2 and 3. UGGT1 adds-back glucose 3, which engages CNX. CST inhibits GI and GII. **B)** CLSM analyses in WT MEF. NS_PL-HA (red) in LAMP1-positive endolysosomes (cyan). Scale bars 10 µm. **C)** Same as (B) in MEF treated with BafA1. **D)** Same as (B) in MEF treated with CST. **E)** Same as (B) in UGGT1 KO MEF. **F)** Same as (B) in CNX KO MEF **G)** LysoQuant [28] quantification of endolysosomes containing the ERLAD client in (A-E). N=3. Mean, n = 7, 15, 16, 29, 21 cells. Welch’s T test of One-way ANOVA, ns P > 0.05, ****P < 0.0001.

### The N-glycan at position 321 in NS_PL signals selection for ERLAD

NS_PL harbors three predicted N-glycosylation sites (N157, N321 and N401). Only those at position N157 and N321 are covalently modified with a N-linked oligosaccharide (**Fig. 4A**) [22]. To confirm this, we performed partial digestion of NS_PL oligosaccharides with two different glycanases, EndoH (**Fig. 4B**, upper panel) and Peptide-N-Glycosidase F (PNGaseF) (**Fig. 4B**, lower panel) on cells transfected with NS_PL. Western blot analysis revealed, with both assays, three distinct bands for NS (**Fig. 4B**, lanes 1-2) and NS_PL (**Fig. 4B**, lanes 3-4) confirming that the proteins carry two (arrows 2NGly in the upper and lower panels, **Fig. 4B**), one (arrows 1NGly), or no glycans (arrows 0NGly). Next, to identify if one of the two NS_PL N-linked oligosaccharides is involved in selection of the folding-defective polypeptide for ERLAD, we generated the glycosylation variants NS_PLN157Q, NS_PLN321Q and NS_PLN401Q (**Fig. 4A**) by replacing the asparagine (N) residue of the N-glycosylation sequons with a glutamine (Q) residue, thereby abolishing N-linked glycosylation. The partial EndoH and PNGaseF digestions reveal only two polypeptide bands for NS_PLN157Q Q (**Fig. 4B**, lanes 5-6, arrows 1NGly and 0NGly) and NS_PLN321Q (**Fig. 4B**, lanes 7-8) confirming that these glycosylation mutants carry only 1 N-linked oligosaccharide. The partial EndoH and PNGaseF digestions reveal 3 polypeptide bands for NS_PLN401Q showing that, as the NS_PL, this form carries 2-linked oligosaccharides (lanes 9-10). These results confirm that NS_PL is glycosylated at positions N157 and N321 (**Fig. 4A**, green ellipses), whereas the N401 is a cryptic glycosylation site (**Fig. 4A**, grey ellipses).

**Fig. 4.**
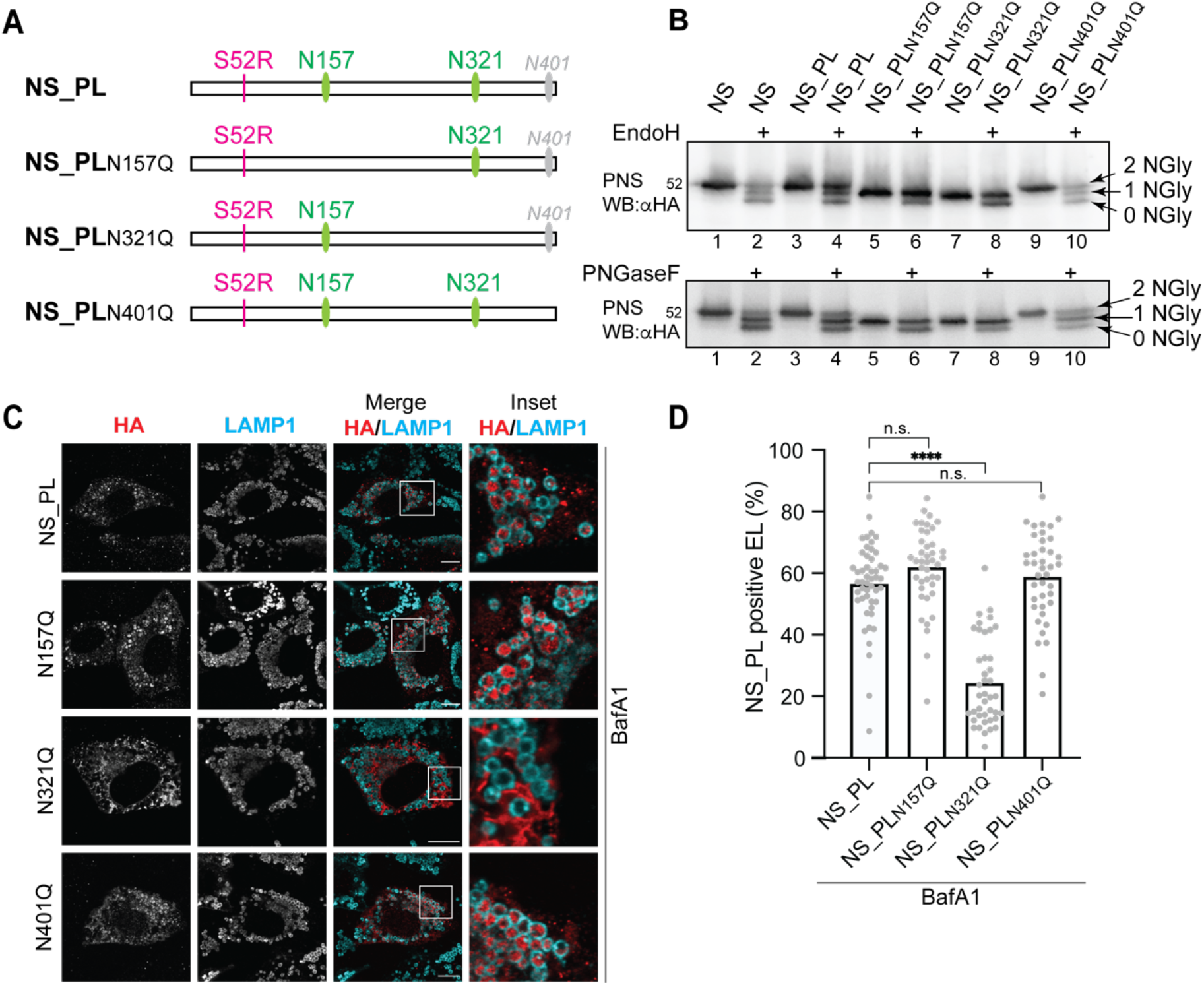
N-glycosylation at position 321 is essential for NS_PL delivery to endolysosomes. **A)** NS_PL and its glycosylation site mutans. Glycosylation sites at N157 and N321 highlighted in green, the cryptic glycosylation site 401 is in grey. The S52R mutation of NS_PL is in pink. **B)** Electrophoretic mobility of NS_PL and the glycosylation mutants expressed in HEK293 analyzed by SDS-PAGE and immunoblotting after partial EndoH or PNGaseF treatment (upper and lower panel, respectively). **C)** CLSM analyses of NS_PL-HA and its glycosylation mutants (red) delivery within LAMP1-positive endolysosomes (cyan) in MEF WT cells treated with BafA1. Scale bars 10 µm. **D)** LysoQuant [28] quantification of HA-positive endolysosomes of C. N=3. Mean, n = 38, 49, 42, 41, 40 cells. Welch’s T test of One-way ANOVA, ns P > 0.05, ****P < 0.0001.

To determine which, if any, of the NS_PL glycans functions as a signaling glycan for ERLAD, we transiently transfected MEF treated with BafA1 with the glycosylation mutants. CLSM reveals that the deletion of oligosaccharide at position 321 substantially hampers the delivery of the misfolded polypeptides within LAMP1-positive endolysosomes (**Fig. 4C, 4D**), whereas the deletions of the oligosaccharides at position N157 and the mutation of the cryptic glycosylation site N401 did not significantly alter the delivery of the misfolded polypeptides within endolysosomes (**Fig. 4C, 4D**).

## Discussion

Misfolded proteins that form large aggregates cannot be dislocated across the ER membrane for proteasomal clearance. Rather, they are segregated in ER subdomains that deliver their toxic content within degradative endolysosomes for clearance by mechanistically distinct ERLAD programs [5–7]. In recent years, ERLAD pathways have been described for an increasing number of clients, indicating that ERAD-resistant substrates are cleared through distinct ERLAD mechanisms that depend on specific ERphagy receptors. For example, ATZ polymers and mutant NPC1 are processed through FAM134B-dependent mechanisms, whereas other aggregation-prone proteins such as secretory acinar enzymes or six-repeat islet amyloid polypeptide (6×IAPP) engage CCPG1, and misfolded proinsulin or pro-arginine vasopressin, and proopiomelanocortin (POMC) rely on RTN3L-mediated ERphagy [6].

The processing of N-linked oligosaccharides determines the fate of the associate polypeptide. Sparse and rare reports show that a single N-glycan within multi-glycosylated proteins can execute a dominant role in N-glycan-regulated folding [40], N-glycan-dependent selection for ERAD [41] or N-glycan-dependent selection for ERLAD [13]. Regarding protein folding, Ari Helenius and colleagues showed that only the glycan at position 81 among the seven N-linked oligosaccharides on influenza virus hemagglutinin (HA) is strictly required for HA maturation [40]. Similarly, for ERAD, Spear and Ng found that during clearance of folding defective, mutant forms of the tetra-glycosylated CPY and of the di-glycosylated PrA proteins from the yeast ER, only the glycans at position 479 of CPY and at position 107 of PrA are necessary to direct these misfolded polypeptides into the ERAD pathway [41]. In our previous work, by tracking the fate of misfolded polymers of the tri-glycosylated Z-variant of alpha1-antitrypsin, we demonstrated that the N-linked glycan at position 83 is crucial for the persistent engagement of CNX, which ensures the FAM134B-mediated ERLAD of this disease-causing polypeptide [13].

With this study, we expand the repertoire of known ERLAD substrates by identifying the aggregation-prone FENIB-causing NS_PL as a novel client of the ERLAD pathway. Our results show that NS_PL is delivered within endolysosomes through LC3-dependent vesicular transport, as indicated by its dependence on LC3 lipidation, the ERphagy receptor FAM134B and the SNARE protein STX17. Notably, also for NS_PL, one of the two N-linked oligosaccharides, the one at position 321, controls the clearance program from the ER. In our study a glycan mutational analysis showed that individual N-glycans differentially regulate the fate of NS_PL. These results together with previous publications strengthens the hypothesis that individual glycans function as signaling appendices for protein folding, ERAD or ERLAD [13,40,41]. An interesting question arising from these findings concerns the structural features that make specific N-glycans dominant signals for ERLAD selection. One possibility is that these glycans are positioned within polypeptide regions characterized by structural or dynamic properties, such as flexible loop regions or locally mobile segments. Such environments may increase the accessibility of the glycan to ER-resident modifiers, including glucosidases, glucosyltransferases, and lectin chaperones, thereby facilitating repeated cycles of glucose trimming, re-glycosylation, and CNX engagement.

## Materials and Methods

### Cell culture, transient transfection, and inhibitors

MEF and HEK293, were grown in DMEM supplemented with 10% FBS at 37°C and 5% CO2. Atg7 (WT and KO) and Atg5 (WT and KO) MEF were gifts from M. Komatsu and N. Mizushima. CNX-KO [37–39] and UGGT1-KO MEF [35,36] have been previously described. SEC62-, FAM134B-, STX17-KO MEF cells were engineered using CRISPR/Cas9 system as previously described [12,32]. Transient transfections were performed using JetPrime transfection reagent (PolyPlus) following manufacturer’s protocol. BafilomycinA1 (BafA1, Calbiochem, Invitrogen) was used at 50 nM for 12 h or 100 nM for 6 h. CST (Sigma) and was used at 1 mM.

### CLSM

MEFs were plated on pre-coated Alcian blue (Sigma) glass coverslips (VWR) prior to transient transfection using the jetPRIME reagent following the manufacturer’s protocol. For LysoQuant analysis, cells were treated with 50 nM BafA1 for 17 h. 24 h post-transfection, cells were fixed at room temperature for 20 min in 3.7% paraformaldehyde (Sigma) in PBS. After three washes with PBS coverslips were then incubated for 20 min in saponin permeabilization solution (PS) (10% goat serum, 10 mM HEPES, 15 mM glycine, and 0.05% saponin in PBS) to allow antibody access to intracellular structures. Following permeabilization, cells were incubated with primary antibodies diluted in PS for 90 min. After three washes with PS, cells were incubated with Alexa Fluor–conjugated secondary antibodies diluted 1:300 in PS for 45 min. Finally, cells were rinsed three times with PS and water and mounted onto glass microscope slides (Epredia) using VECTASHIELD (Vector Laboratories) supplemented with or without 4’,6-diamidino-2-phenylindole (DAPI). Confocal images were captured using STELLARIS 5 and STELLARIS 8 microscopes equipped with a Leica HCX PL APO lambda blue 63×/1.40 oil objective and a pinhole set at 1 a.u. Image acquisition was performed with Leica LAS X software, utilizing diode at 405 nm or laser beams at 489, 499, 552, 561, 587 and 653 nm wavelengths for excitation. Fluorescence emissions were collected at the following ranges: 430–490 nm (Alexa Fluor 405), 504–587 nm (Alexa Fluor 488), 592–644 nm (Alexa Fluor 568) and 658–750 nm (Alexa Fluor 646). Image analysis and quantification were performed using LysoQuant and ImageJ 2.16.0/1.54p software64,123. Image post-processing was performed with Adobe Photoshop (v26.2.0).

### Cell lysis, immunoprecipitation, and Western blot

After the respective treatments, cells were washed with ice-cold phosphate-buffered saline (PBS) containing 20 mM N-ethylmaleimide (NEM). The cells were then lysed using either 2% CHAPS (in HEPES-buffered saline (HBS), pH 6.8) or RIPA buffer (1% Triton X-100, 0.1% SDS, 0.5% sodium deoxycholate in HBS, pH 7.4), both supplemented with 20 mM NEM and protease inhibitors (200 mM phenylmethylsulfonyl fluoride, 16.5 mM chymostatin, 23.4 mM leupeptin, 16.6 mM antipain, 14.6 mM pepstatin) for 20 min on ice. Post-nuclear supernatants (PNS) were collected after centrifugation at 10,600 g for 10 min. The PNS was denatured by adding 100 mM dithiothreitol (DTT) and heated for 5 min at 95 °C before being subjected to SDS–polyacrylamide gel electrophoresis (PAGE). Detergent insoluble material was solubilized in 8 M Urea and dissolved in 0.1 M ammonium bicarbonate for further analyses. For immunoprecipitations, PNSs were diluted with lysis buffer and incubated with Protein G beads (1:10 w/v, swollen in PBS) and select antibodies at 4°C. After three washes of the immunoprecipitants with 0.5% CHAPS in HBS pH 7.4, beads were denatured for 5 min at 95°C and subjected to SDS–PAGE. Proteins were transferred to PVDF membranes using the Trans-Blot Turbo Transfer System (Bio-Rad). Native gel electrophoresis was performed after CHAPS lysis. The PNSs was incubated for 15 min at RT in Native Sample Buffer (Bio-Rad) and run on 7.5% native acrylamide gel in Tris/Glycine Buffer (Bio-Rad). Proteins were then transferred onto PVDF membrane using the Trans-Blot Turbo Transfer System (Bio-Rad). Membranes were blocked with 10% (w/v) non-fat dry milk (Bio-Rad) in Tris-buffered saline containing 1% Tween 20 (TBS-T, Sigma-Aldrich) and stained with primary antibodies diluted in TBS-T. Afterward, horseradish peroxidase (HRP)-conjugated protein A or HRP-conjugated secondary antibodies diluted in TBS-T were applied for 45 min. Membranes were developed using Western Bright ECL and signals captured on Fusion FX Edge 18.12 (Vilber). The quantification of western blot bands was performed using Fusion FX Edge 18.12 and ImageJ 2.16.0/1.54p software123. Membrane stripping for probing additional antigens was done using Re-Blot Plus Strong Solution (Millipore) following manufacturer’s instructions.

### EndoH and PNGaseF assays

For full EndoH (NEB) and PNGaseF (biolabs) treatment, proteins from immunoprecipitated samples or from TCE were split into two aliquots and incubated in the presence or absence of 5 mU of EndoH for 3 h at 37°C according to the manufacturer’s protocol. For partial EndoH and PNGaseF assays samples were instead incubated with 2.5 mU of enzyme for 10 min at 37°C. Samples were then analyzed by SDS–PAGE.

## Abbreviations

BafA1: Bafilomycin A1
CLSM: Confocal Laser Scanning Microscopy
CST: Castanospermine
CNX: Calnexin
EndoH: Endoglycanase H
ER: Endoplasmic Reticulum
ERAD: ER Associated Degradation
ERLAD: ER-to-lysosome Associated Degradation
GI: α-glucosidase I
GII: α-glucosidase II
NS: Neuroserpin
NS_PL: Neuroserpin Portland mutation (S52R)
PNGaseF: Peptide-N-Glycosidase F
PNS: Post Nuclear Supernatant
UGGT1: UDP-glucose:glycoprotein glucosyltransferase
STX17: Syntaxin17

## Acknowledgments

We thank the members of Molinari’s laboratory for discussions, and critical reading of the manuscript.

## Funding

Swiss National Science Foundation grants 310030_214903 and 320030-227541 (MM). Alpha1-Foundation Application ID: 1188512.

## Author contributions

Conceptualization: CH, IF, MM

Methodology: CH, IF, MM

Investigation: CH, IF

Visualization: CH, IF

Funding acquisition: MM

Project administration: MM

Supervision: MM

Writing – original draft: CH, MM

Writing – review & editing: CH, IF, MM

## Competing interests

Authors declare that they have no financial or non-financial interests.

## Notes

### Competing Interest Statement

The authors have declared no competing interest.

